# hnRNPA1 regulates early translation to replication switch in SARS-CoV-2 life cycle

**DOI:** 10.1101/2021.07.13.452288

**Authors:** Ram Kumar, Yogesh Chander, Nitin Khandelwal, Himanshu Nagori, Assim Verma, Yash Pal, Baldev R. Gulati, Bhupendra N. Tripathi, Sanjay Barua, Naveen Kumar

## Abstract

Our study suggests that methylation of severe acute respiratory syndrome coronavirus 2 (SARS-CoV-2) RNA is essential for its optimal replication in the target cells. Heterogeneous nuclear ribonucleoprotein A1 (hnRNPA1, an RNA-binding protein) was shown to mediate deposition of N6-methyladenosine (m6A) in internal SARS-CoV-2 RNA. The levels of hNRNPA1 expression and extent of methylation varied, depending on the course of SARS-CoV-2 life cycle. The recruitment of eIF4E (translational initiation factor) facilitated viral RNA translation at 1 hour post infection (1 hpi). However, at 2 hpi, methylation of internal SARS-CoV-2 RNA recruited hNRNPA1 which facilitated viral RNA transcription but resulted in translational repression, a phenomenon contributing in understanding the early translation to replication switch in the viral life cycle. Besides, the abrogation of methylation also produced a defective 5’ cap of viral RNA which failed to interact with eIF4E, thereby resulting in a decreased synthesis of viral proteins. To conclude, methylation of the internal and 5’ cap of SARS-CoV-2 RNA was shown to regulate transcription and translation of SARS-CoV-2 in a time dependent manner.

**IMPORTANCE:** RNA modifications are found in all life forms and have been linked to development, health and diseases. Our study reveals that internal SARS-CoV-2 RNA methylation (m6A) is essential for interaction with hNRNPA1 to effectively synthesize viral genome. Besides, m6A-marked RNA and hRNPA1 interaction was also shown to regulate early translation to replication switch in SARS-CoV-2 life cycle. Blocking SARS-CoV-2 RNA methylation resulted in reduced virus yield, suggesting epitranscriptomic machinery (methylation) facilitates SARS-CoV-2 replication and might represent potential target for new antiviral drugs against COVID-19.

## INTRODUCTION

Gene expression is not determined solely by the sequence information encoded in the individual’s genome but is rather subjected to multiple levels of control both at the DNA and RNA levels. At the DNA (genomic) level, besides promoters and enhancers, the gene expression is regulated by DNA methylation, histone remodelling, alternative histone variant use and deposition of modifications on histone tails, collectively referred to as epigenetic regulation (1). Similarly, covalent modifications also contribute in determining the stability and translation of mRNA and are referred to as epitranscriptomic gene regulation (2). Out of the over 160 different posttranscriptional modifications described so far, most are abundantly present in ribosomal RNA (rRNA) and transfer RNA (tRNA) (3, 4). The messenger RNA (mRNA) also contains at least 13 different chemical modifications (5) which are grouped as cap-adjacent- and internal modifications (5). Internal modifications occur in coding regions, introns as well as 5’ and 3’ untranslated regions (UTRs) of mRNAs. The N^6^-methyladenosine (m^6^A) is the most abundant internal modification of mRNA and long noncoding RNA (lncRNA) in mammalian cells (5). The enzymes that install, remove and bind to mRNA modifications are called as “writers” “erasers” and “readers” respectively. m^6^A is installed by a methyltransferase complex (writers) containing the core catalytic heterodimer [methyltranferase-like protein 3 (METTL3) and METTL14] and a splicing factor WTAP (Wilms tumor 1-associated protein). The writers bind to short consensus sequence motifs in the target mRNA. m^6^A can be reversibly removed by demethylases (erasers) such as FTO (fat mass and obesity-associated protein) and ALKBH5 (alkylation repair homolog protein 5). m^6^A is widely distributed along the mRNA, although it is enriched around the stop codons and at the 3’ untranslated regions (UTRs) (6, 7). Readers, such as YTH-domain family 2 (YTHDF2) and heterogeneous nuclear ribonucleoprotein (hnRNP), directly or indirectly recognise the m6A-marked transcripts and affect various aspects of mRNA metabolism, including RNA localization, splicing, stability (degradation) and translation. Modifications of cap-adjacent nucleotides are deposited to the 5’-ends of RNAs transcribed by RNA polymerase II. Typically, the cap consists of a 7-methylguanosine (m7G) moiety added in a characteristic 5’–5’ triphosphate linkage to the first transcribed nucleotide. The first and second nucleotides adjacent to the cap can be 2’-O-methylated at the ribose (cOMe) in animals, protists and viruses (8). While the m7G is essential for RNA translation and stability (9), the cOMe of mRNA cap is implicated in the innate host antiviral responses (10).

RNA modifications are found in all domains of life viz; animals, plants and their associated pathogens and have been linked to development, health and diseases (5). m^6^A modifications have also been observed in diverse groups of viruses (11–21) including severe acute respiratory syndrome coronavirus 2 (SARS-CoV-2) (22). Depending on the nature of the virus involved, m6A modifications may either support (15–17, 20) or inhibit (18, 19) viral gene expression. A recent study mapped eight m6A sites in SARS-CoV-2 genome (22). However, the precise role of these m6A marks in SARS-CoV-2 life cycle remains unknown. This study provides functional insights into the epitranscriptomic modifications of the SARS-CoV-2 genome.

## RESULTS

### m6A modifications facilitate SARS-CoV-2 replication

We screened a library of small molecule chemical inhibitors and identified DZNep as an inhibitor with anti-SARS-CoV-2 activity. DZNep is known to act as inhibitors of S-adenosylhomocysteine (SAH) hydrolase and, as a result, to deplete cells of S-adenosylmethionine (SAM), the methyl donor used by METTL3 and several other writers. At the non-cytotoxic concentrations **(FIG S1A),** DZNep exhibited anti-SARS-CoV-2 activity in a dose-dependent manner **(FIG 1A).** Since DZNep did not exert any virucidal effect, anti-SARS-CoV-2 activity of DZNep could be a result of the inhibition of SARS-CoV-2 replication in the target cells, rather than the inactivation of cell free virions **(FIG S1B)**. This suggested that m6A modifications positively regulate SARS-CoV-2 replication. Furthermore, siRNA knockdown of m6A writers (MTTL3) **(FIG S2A),** m6A reader ( hNRNPA1**) (FIG S2B)** and MAT2A-the enzyme involved in synthesizing universal methyl donor (SAM) **(FIG S2C)** resulted in a decreased virus yield **(FIG 1B, 1C and 1D)**, suggesting that m6A modifications are essential for the replication of the SARS-CoV-2 genome.

**FIG 1.**
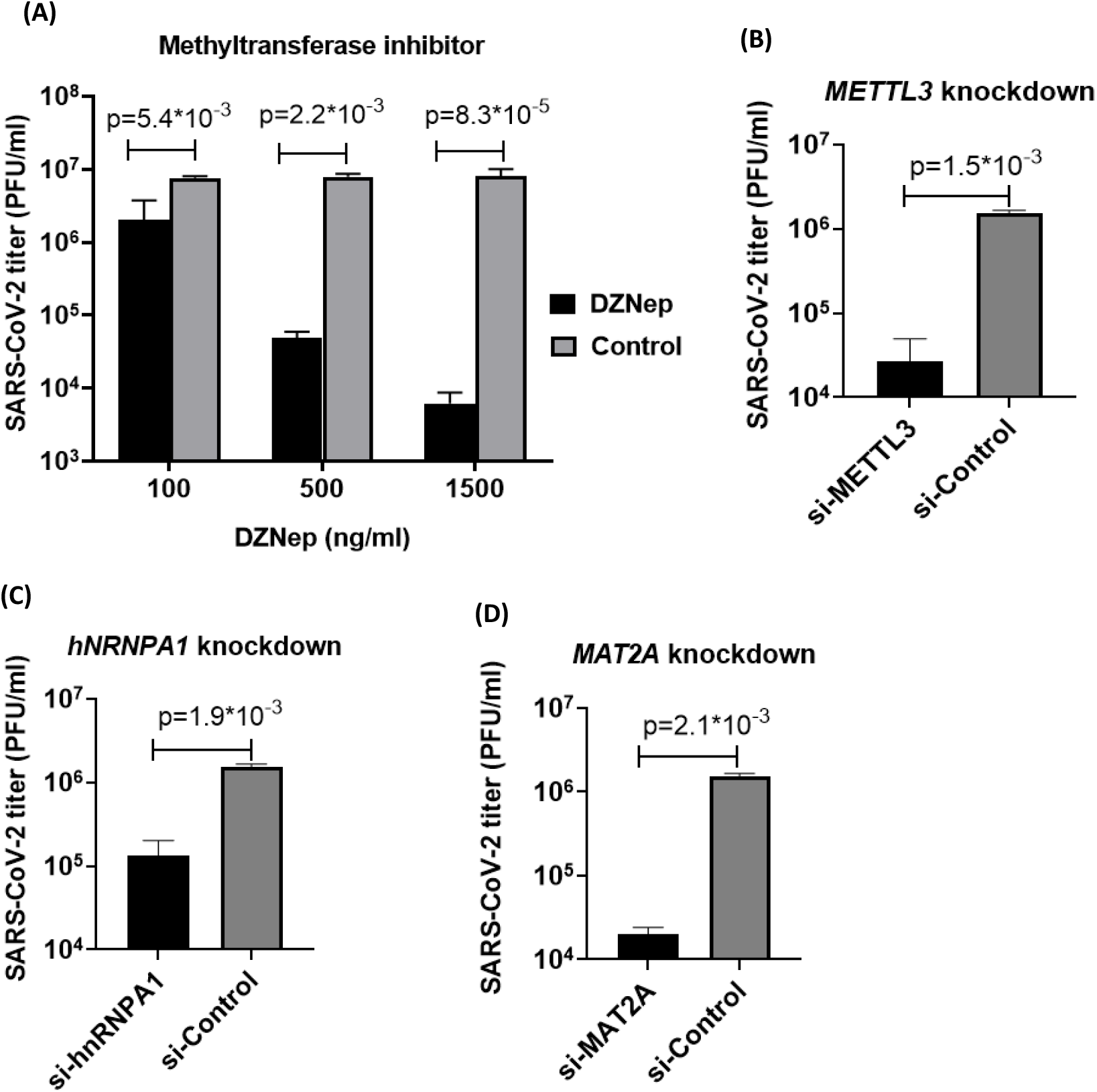
m6A modifications positively regulate SARS-CoV-2 replication. **(A) DZNep inhibits SARS-CoV-2 replication.** Vero cells, in triplicates, were infected with SARS-CoV-2 at MOI of 0.1 in the presence of indicated concentrations of DZNep or vehicle-control. The virus particles released in the infected cell culture supernatants at 48 hpi were quantified by plaque assay. **(B-D) siRNA knockdown.** Vero cells, in triplicates, were transfected with indicated siRNAs along with negative control, followed by SARS-CoV-2 infection at MOI of 1. The virus yields in the infected cell culture supernatants at 24 hpi were quantified by plaque assay. The virus yield in *METTL3* **(B),** *hnRNPA1* **(C)** and *MAT2A* **(D)** knockdown cells is shown. Values are means ± SD and representative of the result of at least 3 independent experiments.

### Reprogramming of m6A methylome in SARS-CoV-2 infected cells

As in other viruses, coronavirus RNA also undergoes epitranscriptomic changes in virus infected cells. A recent study on transcriptome-wide characterization of m6A methylome of SARS-CoV-2 infected cells suggests that m6A sites are widely distributed across the viral RNA (22), although the precise role of these epitranscriptomic marks on the viral genome is yet to be elucidated. We evaluated the kinetics of m6A modifications during the course of SARS-CoV-2 infection in Vero cells wherein cell lysates from SARS-CoV-2-infected cells were subjected to immunoprecipitation using α-m6A, followed by quantitation of viral RNA. We could detect very low (~1%) relative levels of m6A-marked SARS-CoV-2 RNA up to 4 hours post-infection (hpi) **(FIG 2A)** which could not necessarily be due to the low intensity of m6A modifications but rather due to the extremely low levels of total viral RNA at ≤4 hpi, as a significant amount of progeny viral RNA copies were detectable at ~6 hpi in the infected cells (23). However, at 2 hpi, the relative levels of m6A-modified SARS-CoV-2RNA was significantly higher (~1.5%) than at 1 hpi (0.2%) **(FIG 2A)**. The increase in total viral RNA at later time points (≥6 hpi; due synthesis of progeny viral RNA copies), concomitantly resulted in an increase in m6A-marked SARS-CoV-2 RNA as well. The peak levels could be detected at 10 hpi before declining by 12 hpi **(FIG 2A)**. This indicates that SARS-CoV-2 RNA is subjected to m6A modifications in the infected cells and this modification is a dynamic event **(FIG 2A)**.

**FIG 2.**
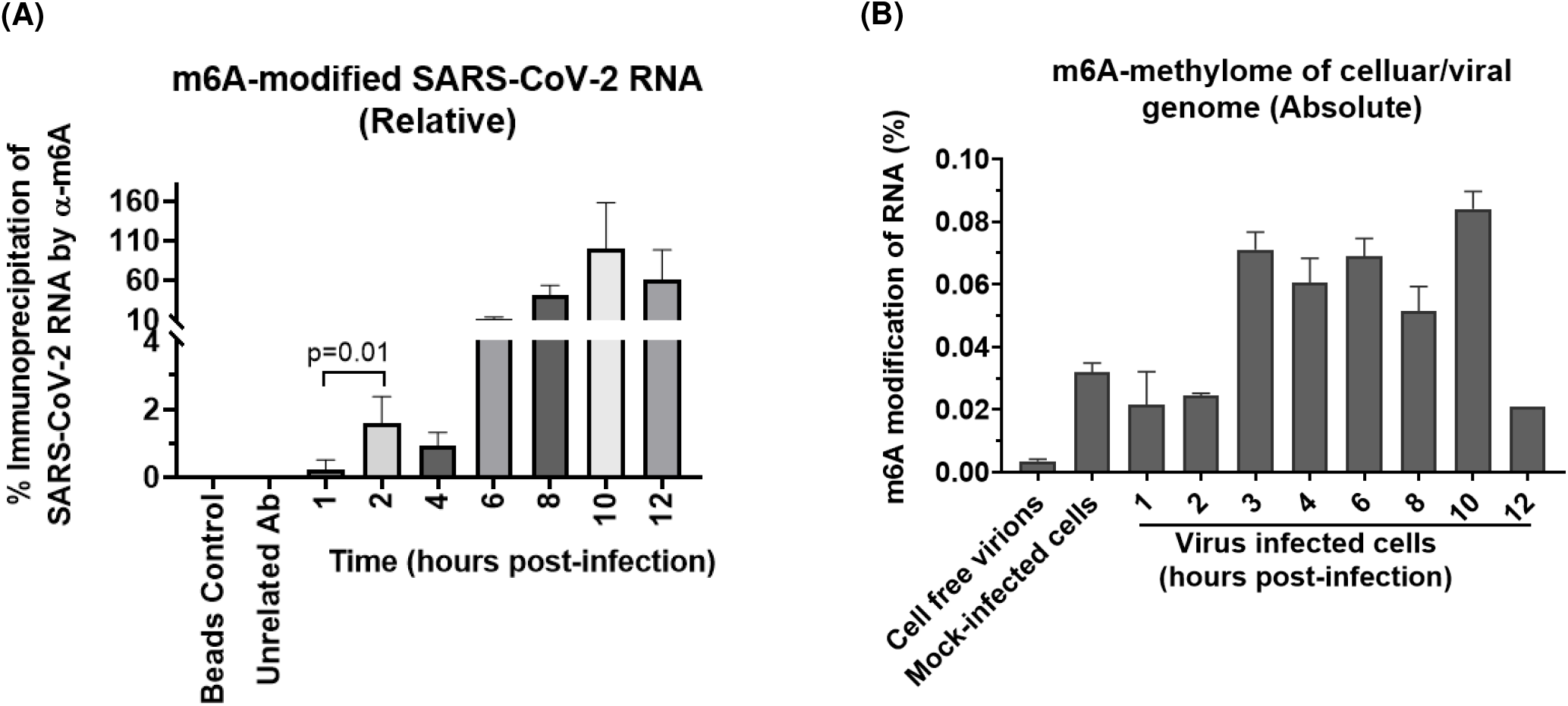
SARS-CoV-2 infection leads to reprogramming of m6A methylome. **(A) Kinetics of the m6A modification in SARS-CoV-2 genome.** Vero cells, in triplicates were infected with SARS-CoV-2 at MOI of 5 followed by washing with PBS and addition of fresh MEM. Cell lysates were prepared at the indicated time points and subjected to CHIP assay. The cell lysates were incubated with α-m6A to immunoprecipitate the m6A-modified RNA. The relative levels of SARS-CoV-2 RNA (“N” gene) in the immunoprecipitate were determined by qRT-PCR. **(B) Quantitation of m6A methylome.** Five hundred millilitre of virus (SARS-CoV-2 infected cell culture supernatant) was filtered using 0.45 μm syringe filter, treated with RNase A and DNAse I to eliminate the uncapsidated cellular RNA/DNA and then ultracentrifuged at 30,000 rpm for 1 h. The resulting pellet was resuspended in 1 ml PBS. Purified virus particles and **c**ell lysates from mock- or SARS-CoV-2-infected cells at indicated time points were subjected to RNA isolation. Equal amount of RNA was evaluated for the determination of the absolute level of m6A modified RNA by EpiQuikTM m6A RNA Methylation Quantification Kit (Colorimetric). The OD values were normalized with negative controls and absolute amount of m6A modified RNA (%) was calculated by comparing it with the positive control. Values are means ± SD and representative of the result of at least 3 independent experiments.

In addition to determining the relative levels of m6A-marked RNA during the course of SARS-CoV-2 replication cycle, we also examined the absolute levels (ratio of m6A-modified RNA versus total RNA) of m6A-marked RNA by EpiQuik m6A RNA Methylation Quantification Kit. In non-infected (mock-infected) cells ~.03% of the total cellular RNA was found to be methylated whereas in SARS-CoV-2-infected cells it varied from 0.02-0.09%, depending on the stage of the virus replication cycle involved-the highest being at middle-late step of virus replication cycle (3 hpi to 10 hpi) and lowest during the initial hours of infection (1 hpi and 2 hpi) **(FIG 2B)**. No detectable amount of methylation was observed in RNA derived from cell-free virions **(FIG 2B).** Taken together, it could be concluded that SARS-CoV-2 RNA is subjected to m6A modifications in the target cells and these modifications are removed before the formation of mature virion particles.

### m6A modifications support synthesis of SARS-CoV-2 RNA

SARS-CoV-2 life cycle is ~10 h in cultured Vero cells (23). In order to evaluate which specific step(s) of SARS-CoV-2 life cycle are affected, DZNep was applied at various times post-infection and the progeny virus particles released at 16 h were quantified. Addition of DZNep had similar levels of suppression in virus yield whether it was applied at 30 min before- or 1 h after virus infection (**FIG S3)**, suggesting that DZNep had no effect on the early steps of SARS-CoV-2 replication. Likewise, there was no inhibitory effect if it was applied at later time points (≥ 6 hpi) of SARS-CoV-2 life cycle which suggested that DZNep does not target the SARS-CoV-2 budding/release. The magnitude of the inhibitory effect of DZNep on SARS-CoV-2 decreased from 1 hpi to 6 hpi, suggesting that DZNep most likely targeted the middle stages (replication/transcription/translation) of SARS-CoV-2 replication. In order to further confirm the specific steps of SARS-CoV-2 targeted by the inhibitors, we conducted the virus step-specific assays (23). The DZNep did not affect the SARS-CoV-2 attachment **(FIG S4A)**, entry **(FIG S4B)** and budding **(FIG S4C).** To further evaluate the effect of the DZNep on the synthesis of viral genome, it was applied at 3 hpi (a time point when early steps of viral life cycle such as attachment and entry are expected to occur) and the cell lysates prepared at 10 hpi, a time point when virus is close to completing its life cycle. DZNep-treated cells exhibited significantly low levels of mRNA **(FIG 3A1)** and total RNA **(FIG 3A2),** suggesting that the m6A modifications could be essential for the optimal synthesis of SARS-CoV-2 genome. In order to confirm the association of m6A marks in synthesizing viral RNA, the cell lysates were immunoprecipitated by α-m6A and quantified by qRT-PCR. The amount of SARS-CoV-2 RNA immunoprecipitated by α-m6A was significantly low in the cells treated with DZNep **[FIG 3B1** (mRNA) and **3B2** (Total RNA)] suggesting that m6A-modifications of SARS-CoV-2 RNA are essential for the optimal synthesis of the SARS-CoV-2 genome.

**FIG 3.**
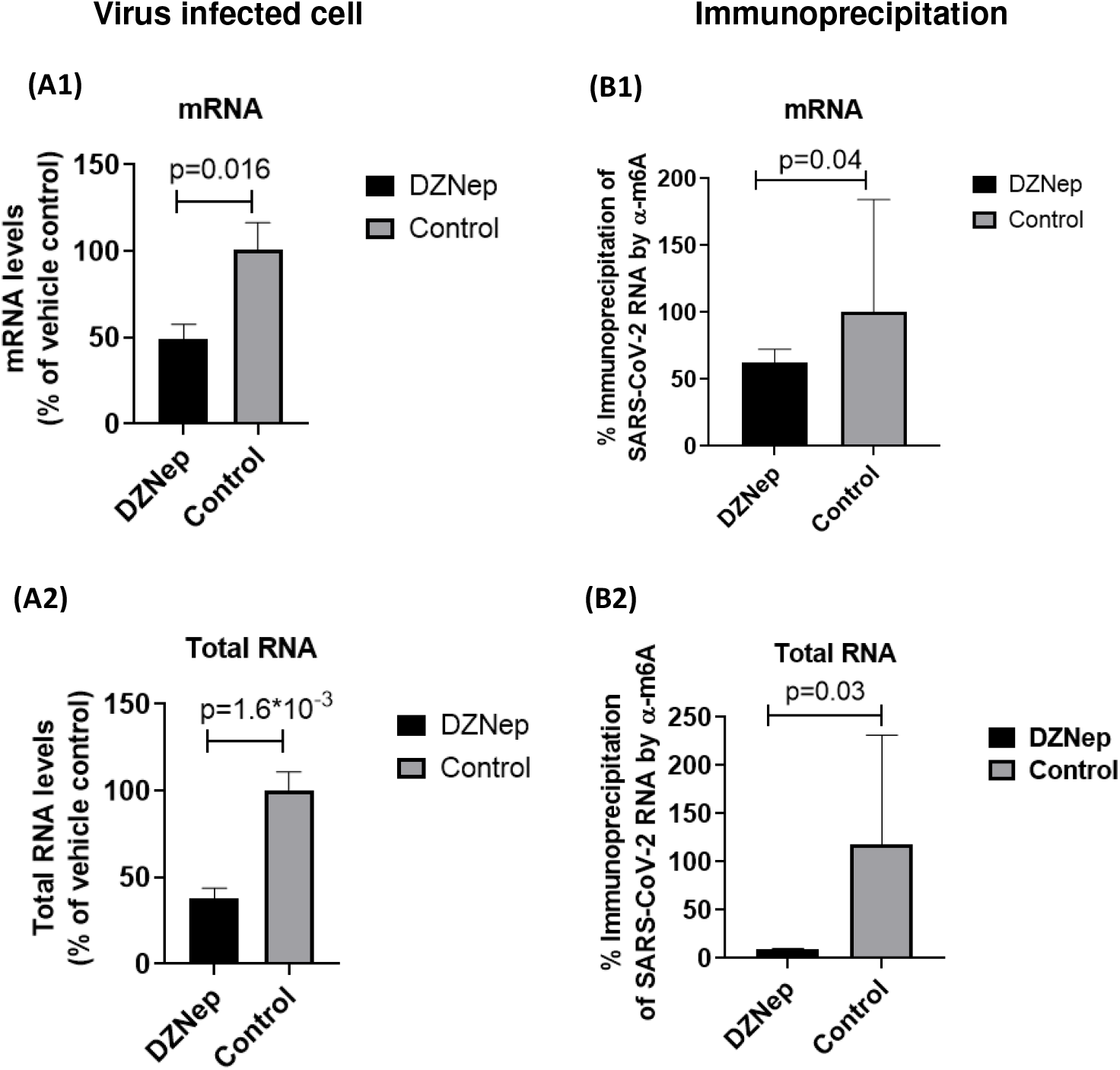
m6A modifications facilitate synthesis of SARS-CoV-2 genome. **(A) Effect of DZNep on synthesis of viral RNA:** Confluent monolayers of Vero cells, in triplicates, were infected with SARS-CoV-2 for 1 h at MOI of 5. DZNep was added at 3 hpi and cells were harvested at 10 hpi to determine levels of SARS-CoV-2 RNA by qRT-PCR. Threshold cycle (Ct) values were analyzed to determine relative fold-change in copy numbers of mRNA **(A1)** and total RNA (**A2**). **(B) m6A modifications are essential for synthesis of SARS-CoV-2 genome.** Vero cells, in triplicates were infected with SARS-CoV-2 at MOI of 5 followed by washing with PBS and addition of fresh MEM. DZNep or vehicle control(s) were applied at 3 hpi and cell lysates were prepared at 8 hpi, Cell lysates were incubated with α-m6A to immunoprecipitate the m6A-modified RNA. The amount of SARS-CoV-2 RNA in the immunoprecipitate was quantified by qRT-PCR. Threshold cycle (Ct) values were analyzed to determine relative fold-change in copy numbers of mRNA **(B1)** and total RNA (**B2**). Values are means ± SD and representative of the result of at least 3 independent experiments.

### hnRNPA1-mediates installation of m6A modifications on SARS-CoV-2 genome

A previous SARS-CoV-2 study has mapped eight m6A sites in the viral genome (22). In order to predict the putative m6A writers (RNA binding proteins), we analysed the SARS-CoV-2 genome (SARS-CoV-2/Human-tc/India/2020/Hisar-4907, bearing Accession Number VTCCAVA294 and GenBank Accession number MW555598) in the RNA-binding protein database (RBPDB) (http://rbpdb.ccbr.utoronto.ca/). Out of the 11 m6A readers known (24), *in silico* binding with the SARS-CoV-2 genome could be predicted with only hnRNPA1 and YTHDC1. The hnRNPA1 was predicted to bind in the 3’-end (two sites) and “S” gene (one site) of the SARS-CoV-2 genome with a position weight matrix score of >9.89 **(Table S1)**. The YTHDC1 was predicted to interact at multiple sites in each gene (except ORF6, ORF7a and ORF7 where no binding sites could be predicted) of SARS-CoV-2, although the position weight matrix score for all the sites was low (<6.5). Next we evaluated the functional role of hnRNPA1 in the SARS-CoV-2 life cycle wherein the SARS-CoV-2 infection in Vero cells resulted in a biphasic expression of hnRNPA1 **(FIG 4A)**. The first and second peak could be observed at 2-4 hpi and 6-8 hpi respectively **(FIG 4A)**. The CHIP assay using α-hnRNPA1 immunoprecipitated the SARS-CoV-2 RNA, suggesting it’s direct interaction with the SARS-CoV-2 RNA **(FIG 4B)**. The DZNep-treatment resulted in a decrease in the SARS-CoV-2 RNA immunoprecipitated by α-hnRNPA1 **(FIG 4B)** which suggested that hnRNPA1-mediated installation of m6A modifications on the viral RNA are essential for optimal synthesis of the viral genome. Further, DZNep did not directly affect the levels of hnRNPA1 expression **(FIG 4C)** suggesting that the inhibitory effect of DZNep is mediated via the RNA-protein interaction rather than by the reduced hnRNPA1 expression.

**FIG 4.**
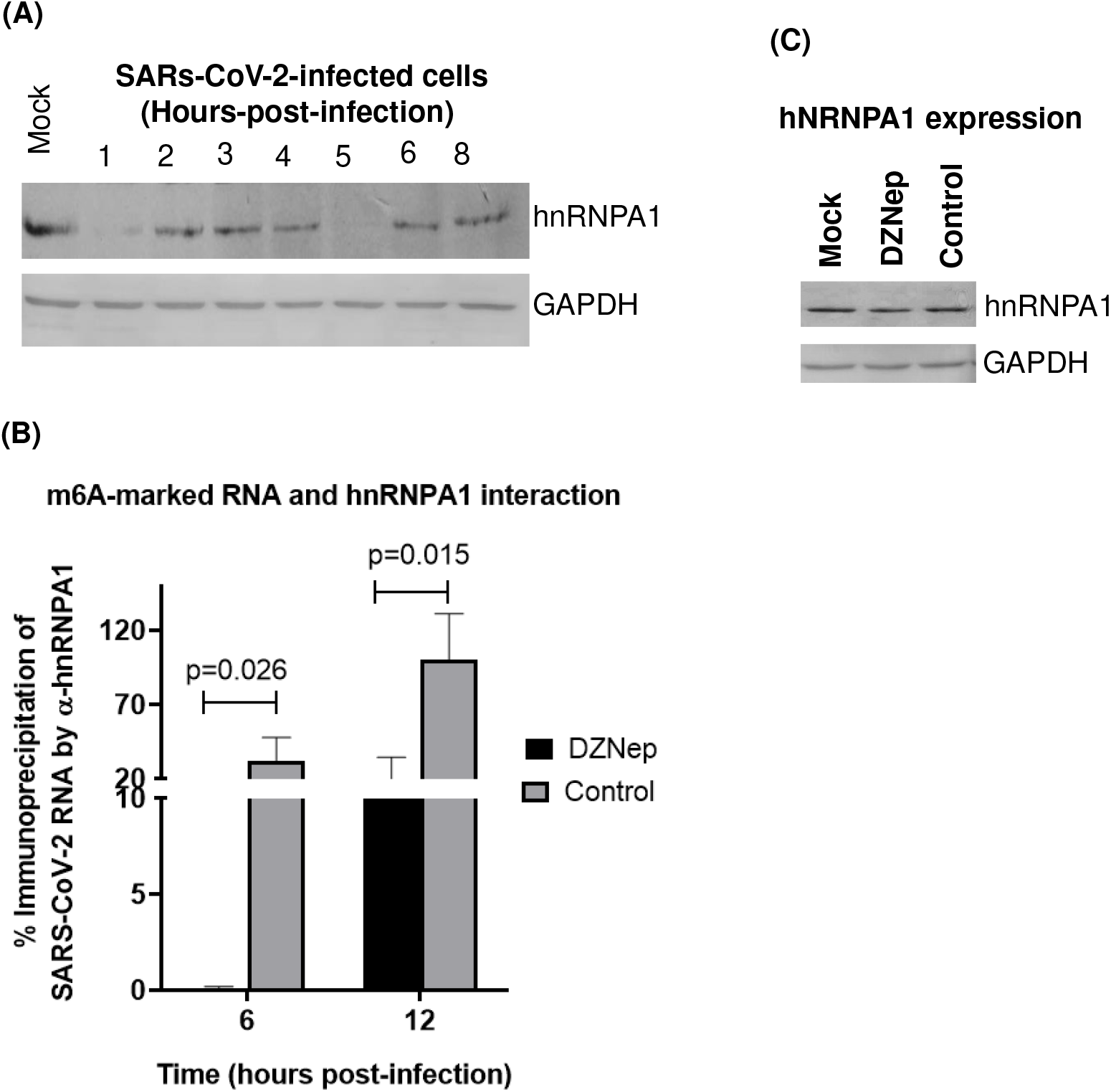
hnRNPA1-mediated installation of m6A modifications facilitate SARS-CoV-2 RNA synthesis. **(A) Kinetics of hnRNPA1 expression in SARS-CoV-2-infected Vero cells.** Vero cells were infected with SARS-CoV-2 at MOI of 5 and the cell lysates were prepared at the indicated time points to determine the levels of hnRNPA1 and GAPDH in a Western blot analysis. **(B) hnRNPA1 interacts with m6A-modified SARs-CoV-2 RNA and this interaction is essential for optimal synthesis of the viral genome.** Vero cells, in triplicates were infected with SARS-CoV-2 at MOI of 5 followed by washing with PBS and addition of fresh MEM. DZNep or vehicle control(s) were applied at 3 hpi and cell lysates were prepared at 6 hpi and 12 hpi, Cell lysates were incubated with α-hnRNPA1 to immunoprecipitate the RNA associated with it. The relative levels of SARS-CoV-2 RNA (E gene) in the immunoprecipitate were determined by qRT-PCR. **(C) Effect of DZNep on hnRNPA1 expression.** Vero cells were infected with SARS-CoV-2 at MOI of 5 and the cell lysates were prepared at 3 hpi to determine the levels of hnRNPA1 and GAPDH in a Western blot analysis. Values are means ± SD and representative of the result of at least 3 independent experiments.

### Methylation of the 5’ cap of SARS-CoV-2 mRNA is essential for eIF4E-mediated translation of viral proteins

Besides inhibiting the RNA synthesis, DZNep treatment also resulted in reduced levels of SARS-CoV-2 proteins in the infected cells (**FIG 5A).** In coronaviruses, viral mRNA translation takes place in a cap-dependent manner wherein the eIF4E plays a central role in the initiation of translation (25–28). Upon phosphorylation by the upstream kinase(s), elF4E binds to the 5’ cap of mRNA to initiate translation (28). Besides internal RNA modifications, the 5’-cap of coronaviruses also undergoes epitranscriptomic changes which include m7G and cOMe (22, 29). The SAM serves as a methyl donor in both these reactions. We further explored if DZNep-induced perturbation of epitranscriptomic changes may affect interaction of the viral mRNA and eIF4E. At 8 hpi (when SARS-CoV-2 RNA was expected to be at its peak level), cells were covalently cross-linked and evaluated for viral RNA and eIF4E interaction in a CHIP assay. In agreement with our previous findings (23), α-eIF4E (reactive antibody) but not α-ERK (non-reactive antibody) or beads control immunoprecipitated SARS-CoV-2 RNA (**FIG 5B).** The levels of viral RNA immunoprecipitated by α-eIF4E were ~99.9% lower in DZNep-treated cells as compared to the vehicle-control-treated cells (**FIG 5B)** which suggested that DZNep inhibits eIF4E/SARS-CoV-2 mRNA interaction which may eventually result in the decreased synthesis of viral protein. In qRT-PCR, the levels of SARS-CoV-2 RNA in α-ERK-treated cells (but not α-eIF4E-treated cells) were undetectable which clearly indicated that α-eIF4E specifically interacted with SARS-CoV-2 RNA (**FIG 5B**).

**FIG 5.**
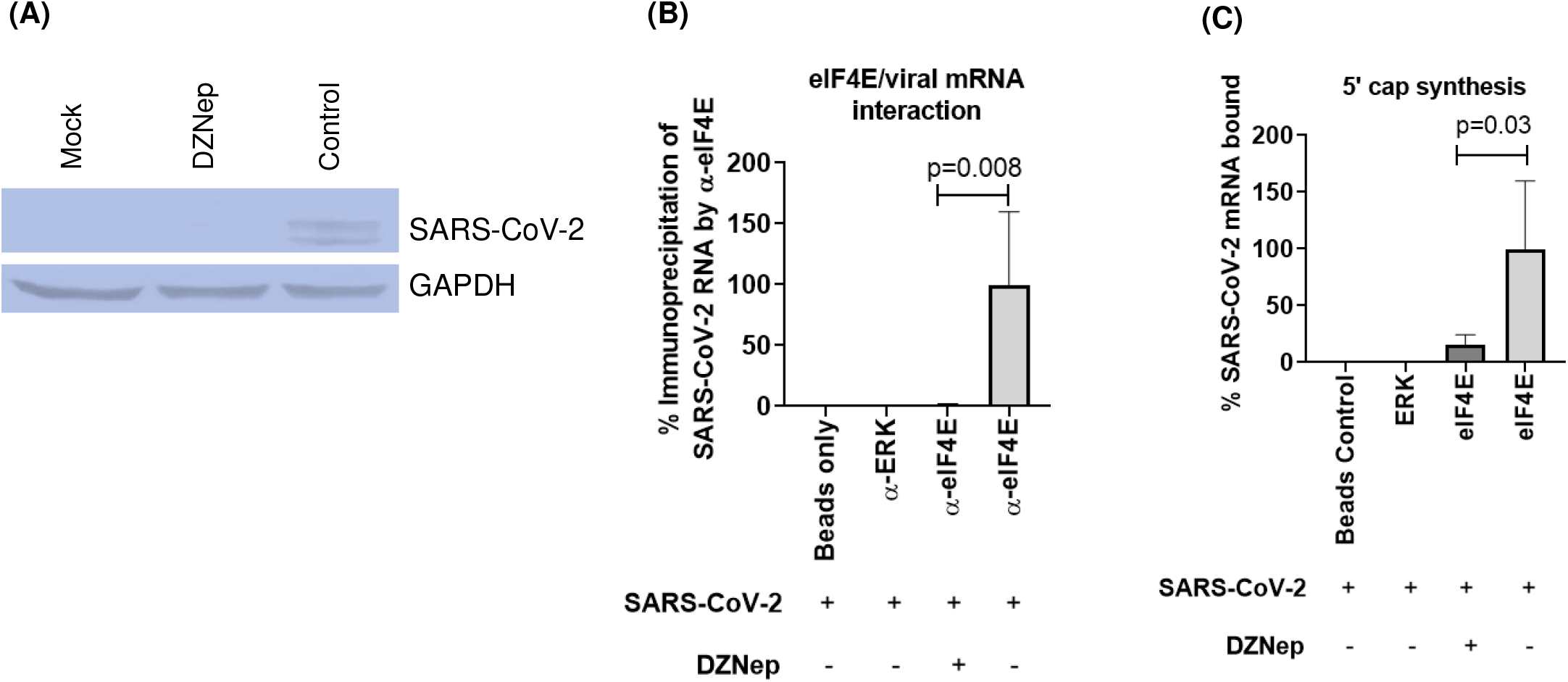
Methylation of the cap-adjacent nucleotides in the 5’ cap of SARS-CoV-2 mRNA is essential for eIF4E-mediated translation of viral proteins. **(A) Effect of DZNep on synthesis of SARS-CoV-2 protein.** Vero cells were infected with SARS-CoV-2 at MOI of 5. DZNep or DMSO were added at 3 hpi. Cell lysates were prepared at 8 hpi to detect the levels of viral proteins by Western blot analysis. The levels of viral proteins **(upper panel)**, along with housekeeping GAPDH protein (**lower panels**) are shown. **(B) Cap adjacent epitranscriptomic modifications in the 5’ cap of SARS-CoV-2 mRNA are essential for interaction with eIF4E.** Vero cells, in triplicates were infected with SARS-CoV-2 at MOI of 5 followed by washing with PBS and addition of fresh MEM. DZNep or vehicle control(s) were applied at 4 hpi and cell lysates were prepared at 8 hpi, Cell lysates were incubated with α-eIF4E to immunoprecipitate the RNA. The amount of SARS-CoV-2 RNA in the immunoprecipitate was quantified by qRT-PCR. Threshold cycle (Ct) values were analyzed to determine relative fold-change in copy numbers of SARS-CoV-2 mRNA (N gene). **(C) DZNep leads to defective synthesis of the 5’ cap of viral mRNA.** The p-eIF4E was purified from uninfected Vero cells as described in the material and method section. Next, Vero cells, in triplicates were infected with SARS-CoV-2 at MOI of 5 followed by washing with PBS and addition of fresh MEM. DZNep or vehicle control(s) were applied at 3 hpi. At 8 hpi, cells were subjected to RNA extraction using TRI reagent. Equal amount of viral RNA from DZNep and vehicle control-treated cells (RNA levels were normalized by qRT-PCR) were incubated with purified peIF4E (described above) for 30 minutes. The immunoprecipitate was subjected to RNA extraction, cDNA preparation (using oligo dT) and quantitation of SARS-CoV-2 mRNA (N gene) by qRT-PCR. Values are means ± SD and representative of the result of at least 3 independent experiments.

Besides inhibiting the interaction between SARS-CoV-2 mRNA and eIF4E, alternatively, the DZNep-mediated decreased synthesis of viral proteins could also be due to defective synthesis of the 5’ cap of viral mRNA. We performed a cell-free viral mRNA and cellular phospho-eIF4E (p-eIF4E) interaction assay wherein p-eIF4E (purified from the mock-infected cells lysate using α-peIF4E) was subjected to *in vitro* interaction with SARS-CoV-2 RNA isolated either. Since the reduce amount of viral RNA in DZNep-treated cells could eventually reflect lower levels of immunoprecipitated viral RNA, equal amount of viral RNA from vehicle control-and DZNep-treated cells (normalized by qRT-PCR) was taken for the assay. As compared to the vehicle control-treated cells, the amount of RNA immunoprecipitated by α-peIF4E-peIF4E complex was ~80% lower in cell lysates from DZNep-treated cells (**FIG 5C)**. This confirmed that perturbation of epitranscriptomic machinery (methylation) produce defective SARS-CoV-2 mRNA which is unable to interact with eIF4E, eventually resulting in decreased synthesis of viral proteins (23).

### m6A modifications regulate the switching from translation to replication of SARS-CoV-2 RNA

SARS-CoV-2 has a positive-sense RNA genome. Immediately upon infection, 2/3^rd^ of the nascent viral RNA is directly translated into 16 non-structural proteins (NSPs) in the cytoplasm of the infected cells (30). The same nascent viral RNA then switches to act as template for viral RNA transcription. We observed a switch on/off phenomenon in the hRNPA1 expression levels in SARS-CoV-2 infected Vero cells **(FIG 4A).** This tempted us to speculate that hRNPA1 could be involved in switching from translation to replication of SARS-CoV-2 RNA. At 1 hpi, we could detect low levels of hNRNPA1 (m6A reader) **(FIG 4A**) and viral/cellular m6A methylome **(FIG 2A** and **2B)** but higher levels of p-eIF4E- the cellular protein that participates in translation) **(Fig 6a)**. Conversely, at 2 hpi, the levels of methylome **(FIG 2A** and **2B)** and hNRNPA1 **(FIG 3A)** were significantly higher but the level of p-eIF4E decreased **(FIG 6A)**. Differential recruitment of translational (p-eIF4E) and transcriptional cellular machinery (m6A/hNRNPA1) respectively at 1 hpi and 2 hpi indicated that these cellular factors may mediate the switching between translation and transcription. Since the recruitment of these epitranscriptomically-regulated cellular factors was dampened in the presence of DZNep, the switching phenomenon could be associated with epitranscriptomic changes of viral mRNA. To provide further insights into this switching phenomenon, SARS-CoV-2-infected Vero cells were cross-linked at 2 hpi and evaluated for the interaction of m6A-modified RNA with RNA-binding proteins (p-eIF4E/hnRNPA1) in CLIP assay. The hnRNPA1 principally binds at the 3’-end of the SARS-CoV-2 genome **(Table S1)** whereas the eIF4E is a 5’ mRNA cap-binding protein. Therefore, the sonication step was omitted in the CLIP assay which ensured the recovery of all the RBPs associated with SARS-CoV-2 genome. The cross-linked m6A-modified RNA was immunoprecipitated by α-m6A and the associated RBPs viz; hnRNPA1 and p-eIF4E were probed in Western blot analysis. At 2 hpi, only hnRNPA1 but not eIF4E was shown to be associated with m6A-modified SARS-CoV-2 RNA, suggesting repression of translation (2 hpi) due to recruitment of hnRNPA1 (**FIG 6B)**. The amount of hNRNPA1 immunoprecipitated by α-m6A was lower in DZNep-treated cells as compared to the control **(FIG 6B)** which suggested a decreased synthesis of SARS-CoV-2 genome due to perturbation of the epitranscriptomic machinery.

**FIG 6.**
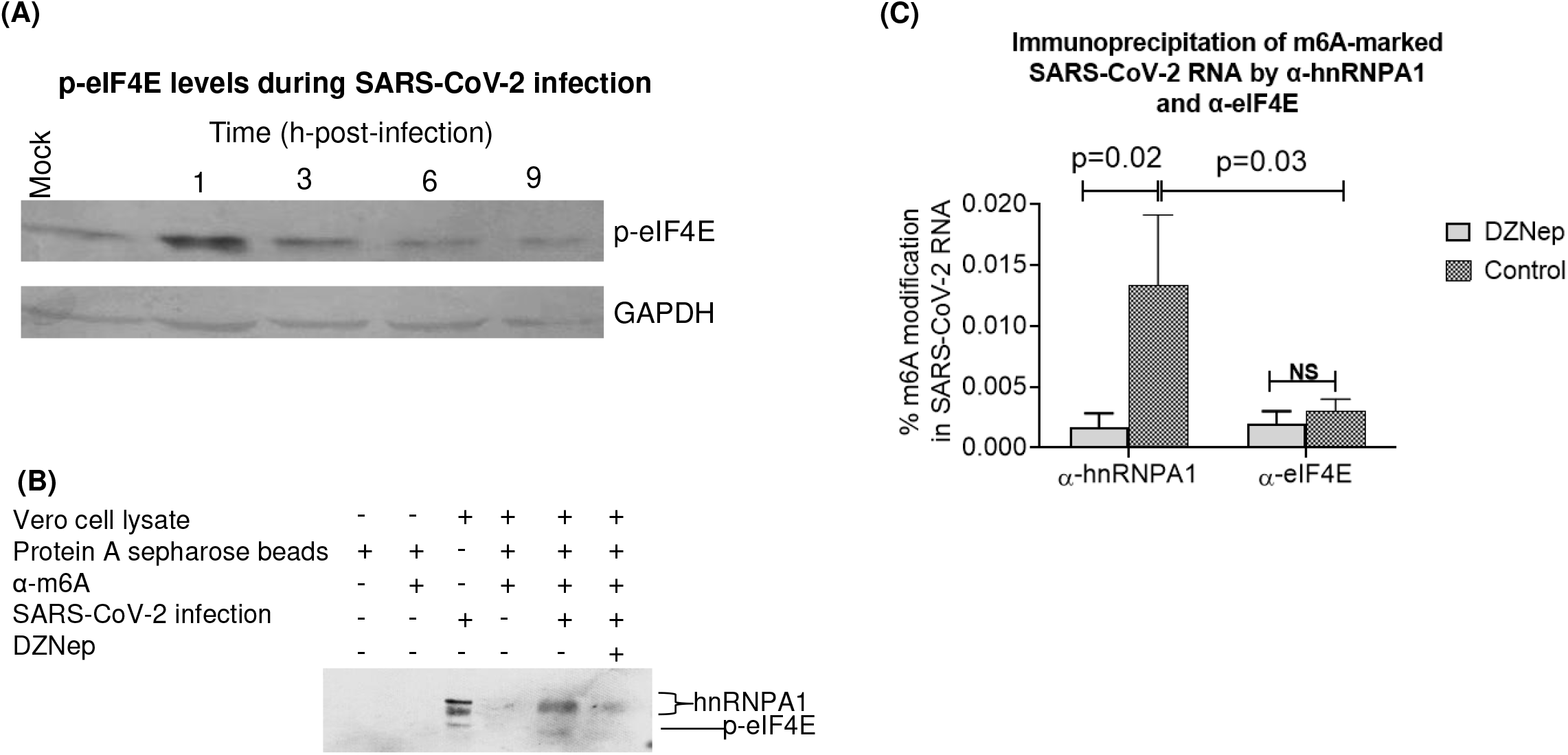
m6A modifications of viral RNA act as a molecular switch from translation to replication of SARS-CoV-2 RNA. **(A) Kinetics of eIF4E activation in SARS-CoV-2-infected cells.** Vero cells were infected with SARS-CoV-2 at MOI of 5 and the cell lysates were subjected to determination of the levels of p-eIF4E and GAPDH in a Western blot analysis. **(B) Levels of α-hnRNPA1 and α-eIF4E in the cell lysate immunoprecipitated by α-m6A**. Confluent monolayers of Vero cells were infected with SARS-CoV-2 at MOI of 5 for 1 h in the presence of DZNep or vehicle control, followed by washing with PBS and the addition of fresh MEM having either DZNep or vehicle control. At 2 hpi cells were subjected to covalently cross-link proteins and nucleic acid for 10 min. The cells lysates and cytosolic fractions were prepared as described in materials and methods under CLIP assay. The cytosolic fraction was subjected to immunoprecipitation by α-m6A. Proteins (hnRNPA1 and eIF4E) interacting with m6A-marked-RNA were probed from the immunoprecipiate (protein-RNA complex) by Western blot analysis. **(C) Levels of m6A-modifed SARS-CoV-2 RNA in the cell lysate immunoprecipitated with α-hnRNPA1 and α-eIF4E.** Confluent monolayers of Vero cells were infected with SARS-CoV-2 at MOI of 5 for 1h in the presence of DZNep or vehicle control, followed by washing with PBS and the addition of fresh MEM having either DZNep or vehicle control. At 2 hpi, cells were subjected to covalent cross-linking. The cells lysates were then incubated with α-hnRNPA1 or α-eIF4E and the immunoprecipiate was probed for determination of m6A methylome by EpiQuikTM m6A RNA Methylation Quantification Kit (Colorimetric). Values are means ± SD and representative of the result of at least 3 independent experiments.

To further confirm the association of m6A modifications in regulating the switching from translation to replication, we performed another assay wherein SARS-CoV-2 infected cell lysates were subjected to immunoprecipitation by α-hnRNPA1 and- α-pEIF4E, followed by quantitation of m6A RNA methylome in the immunoprecipitates. As shown in **FIG 6C,** the amount of m6A-modified total RNA immunoprecipitated by α-hnRNPA1-was much higher than those immunoprecipitated by α-p-eIF4E, suggesting that hnRNPA1 (but not eIF4E) interacts with m^6^A-marked RNA or in other words m6A marks on SARS-CV-2 allows recruitment of RBPs such as hnRNPA1 which represses translation and facilitate transcription. Further, the amount of m6A-modified RNA immunoprecipitated by α-hnRNPA1 was found to be significantly lower in DZNep-treated cells as compared to the vehicle-control-treated cells, suggesting an arrest in RNA synthesis due to perturbation of epitranscriptomic (m6A modification) machinery. These multiple lines of evidence clearly suggest that m6A modifications of viral RNA serve as a molecular switch from translation to replication during early hours of SARS-CoV-2 replication.

## DISCUSSION

The rapidly emerging field of epitranscriptomics has mapped several modifications in mRNA and their impact on gene expression. Only 2–5% of cellular RNA is mRNA (5). Further, the abundance of mRNA modifications is low, with a prevalence rate of <0.5% of all nucleotides (31, 32). These modifications are variable in the range of 5-88% at a given site or transcript (33, 34). m^6^A is the most predominant epitranscriptomic modification in mRNA (11–19, 22), accounting for 0.1-0.5% of all adenosines (35). Despite an extremely low distribution, they play important roles in cellular homeostasis (32, 36) and are dysregulated in several disease conditions including viral infections (37–39). SARS-CoV-2 infection also triggers a global increase in m6A methylome (22). The levels of m6A methylome in our study varied from 0.02-0.09% **(FIG 3B),** depending on the course of virus infection. Since cell free SARS-CoV-2 particles do not harbour any significant m6A modifications (<0.003%) **(FIG 3B),** these are deposited inside the host cell and removed before formation and budding of the mature virion particles. However, some viruses can maintain a relatively high amount of m6A-modified genome in the intact virion particles (40), suggesting a complex pattern of epitranscriptomic regulation of virus replication.

To gain a detailed view on the impact of methylation on virus replication, we treated the cells with a sub-cytotoxic concentration of DZNep (methyl transferase inhibitor) and examined its effect on the individual steps of SARS-CoV-2 replication. DZNep decreased the levels of viral genome and protein synthesis, without affecting other steps of the viral life cycle such as attachment, entry and budding. The participation of m6A machinery in synthesizing SARS-CoV-2 genome was further confirmed by a reduced immunoprecipitation of SARS-CoV-2 RNA by α-m6A in DZNep treated cells.

Next we identified the reader of m6A modification for SARS-CoV-2 RNA. *In silico* binding studies, immunoprecipitation of SARS-CoV-2 RNA by α-hnRNPA1, together with lower levels of RNA immunoprecipitation in the DZNep treated cells confirmed that hnRNPA1serves as an m6A reader and facilitates the synthesis of SARS-CoV-2 RNA. Since DZNep did not directly affect the level of hnRNPA1 expression, its inhibitory effect is mediated via RNA-protein interaction rather than by reduced hnRNPA1 expression.

Besides low levels of viral RNA, DZNep treatment also resulted in a decreased synthesis of SARS-CoV-2 proteins. This could be a reflection of low levels of mRNA or direct interruption in protein synthesis. In coronaviruses, viral mRNA translation takes place in a cap-dependent manner (41) wherein the eIF4E plays a central role in the initiation of translation (25–28). Upon activation by upstream kinase(s), elF4E binds to the 5’ cap of mRNA to initiate translation (28). The 5’-cap structure of viral mRNA also undergoes at least two epitranscriptomic (methylation) modifications viz; m7G and cOMe (22, 29). A reduced immunoprecipitation of SARS-CoV-2 mRNA by α-peIF4E from DZNep-treated cells suggested that methylation (m7G and cOMe) of the 5’ cap of viral mRNA is essential for its interaction with eIF4E-a prerequisite for translation of viral proteins (23). Alternatively, reduced levels of viral proteins in the DZNep-treated cells could also be due to defective synthesis of the 5’ cap of viral mRNA. In a cell-free viral mRNA and cellular p-eIF4E interaction assay, perturbation of the epitranscriptomic machinery by DZNep resulted in the production of viral RNA which was not able to interact properly with eIF4E, suggesting that methylation of the cap-adjacent nucleotides of SARS-CoV-2 mRNA is essential for the proper formation of the 5’ cap.

Immediately following infection, the nascent positive sense viral RNA is directly translated to produce a polyprotein which is further cleaved to produce 16 non-structural proteins (NSPs), which then facilitates the transcription of genomic and subgenomic RNAs (41, 42). We reveal that this early switch from translation to replication in the viral life cycle is regulated by epitranscriptomic modifications. Initially we observed a switch on/off phenomenon in the hnRNPA1 expression levels in SARS-CoV-2 infected Vero cells **(FIG 4A).** This, together with previous studies on m6A-mediated repression of cellular translation (43), tempted us to speculate that hnRNPA1 could be involved in switching from translation to replication of SARS-CoV-2 RNA. It has been hypothesized that besides viral proteins, RBPs also play an important role in switching from translation to replication of the viral genome (44–46), although there is very limited experimental proof (45). Recruitment of cellular factors associated with translation (p-eIF4E) at 1 hpi and those associated with transcription (methylome/hNRNPA1) at 2 hpi indicated the involvement of epitranscriptomic marks in switching from translation to transcription. Perturbation of m6A pathway (2 hpi) resulted in a defective hnRNPA1-mediated synthesis of SARS-CoV-2 RNA which further confirmed the recruitment of epitranscriptomic machinery during viral transcription **(FIG 6B)**. Furthermore, we also demonstrated (at 2 hpi) that hnRNPA1 (but not eIF4E) interacts with m^6^A-marked internal RNA and inhibition of m6A modification results in an arrest in hnRNPA1-mediated synthesis of SARS-CoV-2 genome **(FIG 6C).** These lines of evidence clearly suggest that installation of m6A marks in the SARS-CoV-2 RNA recruit hnRNPA1 that eventually facilitates transcription and represses translation **(FIG 7),** a novel role of epitranscriptomic machinery in regulating switch from translation to replication in SARS-CoV-2 life cycle not reported so far.

**FIG 7.**
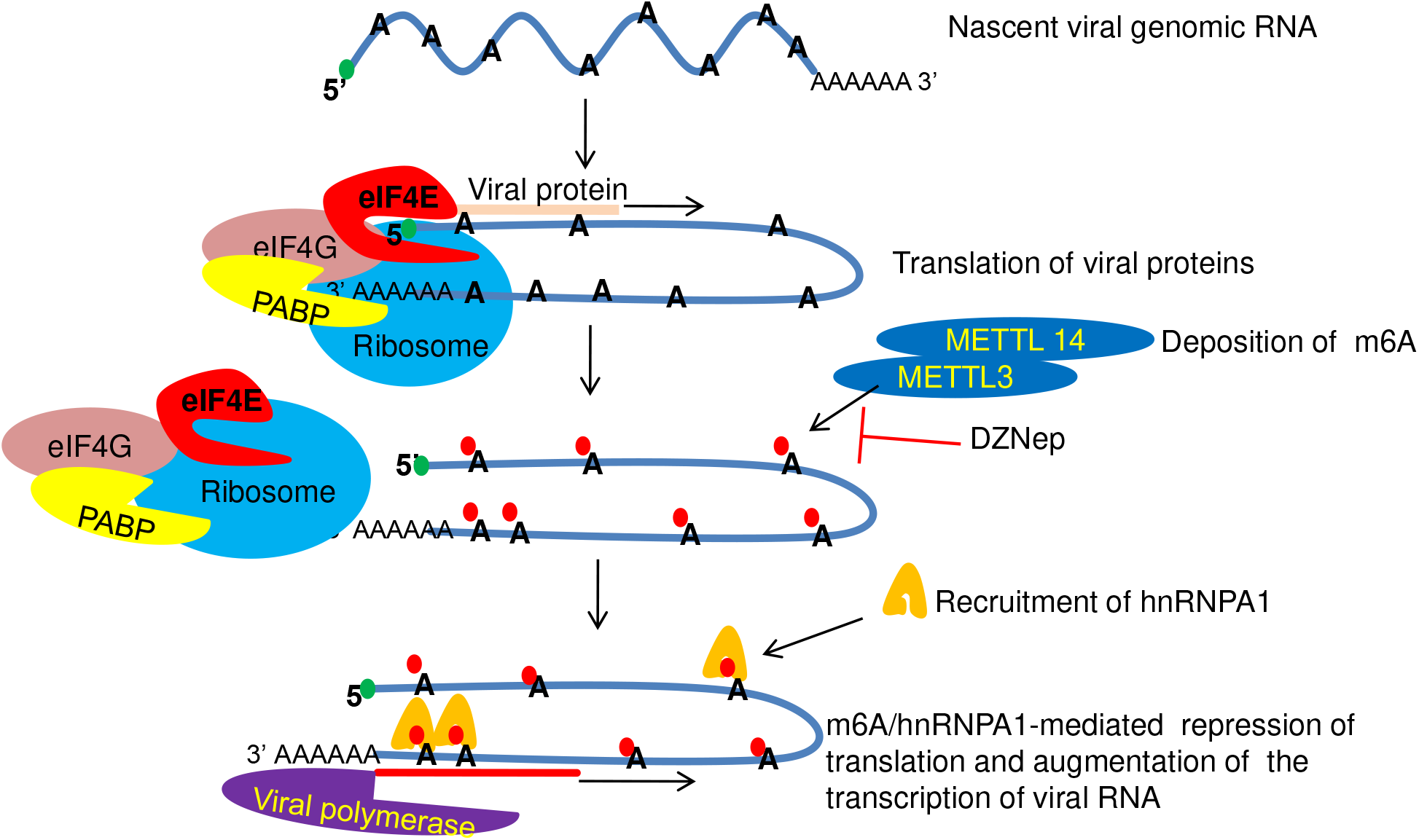
Role of epitranscriptomic machinery in SARS-CoV-2 replication. Immediately following infection (~1h), the nascent positive sense SARS-CoV-2 RNA interacts with cap-dependent translational initiation machinery to directly translate the viral polyprotein which is further cleaved to produce 16 NSPs. After sometime (~2h), viral RNA is subjected to m6A modifications (eight m6A sites in SARs-CoV-2 genome) via cellular writers such as METTL3 and METTL14. m6A deposition facilitates recruitment of hnRNPA1 (three hnRNPA1 binding sites-two at 3’ end and one in “S” gene) which eventually repress translation and facilitate transcription-switch of viral RNA from translation to transcription. DZNep treatment inhibits deposition of m6A mark on SARS-CoV-2 RNA which eventually inhibits recruitment of hnRNPA1 and hence reduced synthesis of the viral RNA.

Methyltransferase inhibitor DZNep together with siRNA knockdown of MTTL3 (m6A writer), hNRNPA1 (m6A reader) and MAT2A (enzyme that participates in the synthesis of SAM-the universal methyl donor) in Vero cells resulted in a reduced virus yield indicating that the m6A epitranscriptomic machinery facilitates SARS-CoV-2 replication and could serve as a novel target for antiviral drug development. Inhibitors that block the m6A pathway would target cellular factors (47), therefore the emergence of drug-resistant viral mutants would seem to be unlikely in epidrugs (28, 48–55). However, since such drugs would also inhibit the epitranscriptomic modification of cellular mRNAs, they may lead to toxicity (48) and should be used for treating the acute infections. Nevertheless, the majority of the host-directed agents which are in clinical use against cardiovascular and inflammatory diseases or cancers have minimal or no adverse side effects (56).

Depending on the nature of the virus replication, depletion of the m6A machinery may have differential pro-(15–17, 20) and anti-viral (18, 19) impact on viral life cycle (57). A recent study by Liu et al., (22), which reveals the negative impact of m6A methylome on SARS-CoV-2 gene expression seems to contradict our findings. However, this study is based on measuring “S” protein expression as an indicator of virus replication, rather than quantifying virus yield. m6A modification is a dynamic event (57, 58). Our study suggests that methylation may differentially regulate transcription and translation of SARS-CoV-2 genome, depending on the course of virus replication.

The effectiveness of the epigenetic machinery may depend on the environment (59, 60), microbiota (61), nutrition (62) and comorbidities (63), implying that a given viral RNA genome may perform differently in different individuals. In this context, our study which highlights the impact of an epitranscriptomic layer of regulation on the life cycle of SARS-CoV-2 is likely to contribute in understanding the pathogenicity, transmission and disease severity in the COVID-19 patients. To conclude, our study reveals that epitranscriptomic modifications of SARS-CoV-2 RNA may differentially regulate transcription and translation of the viral genome. These modifications were substantiated to positively regulate SARS-CoV-2 replication and serve as new pharmaceutical target for antiviral drug development against COVID-19.

## MATERIAL AND METHODS

### Ethics Statement

The study involves collection of biological specimens from humans. Serum samples (1 ml each) were collected from the COVID-19 positive patients who were admitted to Maharaja Agarsen Medical College, Agroha, Hisar, India for treatment. Samples were received via civil surgeon (District Medical Officer) Hisar, India who granted the permission to collect the biological specimens. A due consent was also taken from the patient before collection of the specimens.

### Cells

African green monkey kidney (Vero) cells, available at the National Centre for Veterinary Type Cultures (NCVTC), Hisar were grown in Minimum Essential Medium (MEM) supplemented with 10% fetal bovine serum (FBS) (Sigma, St. Louis, USA) and antibiotics (Penicillin and Streptomycin).

### Virus

SARS-CoV-2/Human-tc/India/2020/Hisar-4907 bearing Accession Number VTCCAVA294 along with its whole genome sequencing data (GenBank Accession number MW555598) has been described previously by our group (23). SARS-CoV-2 was propagated in Vero cells in the Biosafety level 3 (BSL-3) laboratory of ICAR-National Research Centre on Equines (NRCE), Hisar, India. The virus was quantified by plaque assay and viral titers were measured as plaque forming unit per millilitre (PFU/ml) (23).

### Inhibitor

3-Deazaneplanocin A (DZNep) was procured from BioGems International Inc. (Westlake Village, CA, USA) and dissolved in dimethyl sulfoxide (DMSO).

### Antibodies

N6-Methyladenosine (m6A) (D9D9W) Rabbit mAb, METTL3 (E3F2A) Rabbit mAb and hnRNP A1 (D21H11) monoclonal antibodies were received from Cell Signalling Technology (Massachusetts, USA). eIF4E monoclonal antibody (5D11) and Phospho-eIF4E (Ser209) polyclonal antibodies were received from Invitrogen (South San Francisco, CA, USA). Mouse anti-GAPDH (Glyceraldehyde 3-phosphate dehydrogenase, house-keeping control protein) primary antibody, Anti-Mouse IgG (whole molecule)-Alkaline Phosphatase antibody (produced in goat) and Anti-Rabbit IgG (whole molecule)–Peroxidase antibody (produced in goat) was received from Sigma-Aldrich (St. Louis, USA). Rabbit anti-human IgG–HRP was procured from GeNei™, Peenya (Bangalore, India). Human serum from a COVID-19 confirmed patient was collected at 2 weeks following recovery and obtained after due consent from the patient.

### Determination of cytotoxicity and virucidal activity of DZNep

The cytotoxicity of chemical inhibitors in Vero cells was determined by MTT assay as described previously by our group (64).

### Quantitative real-time PCR

Until specified, viral RNA was extracted by QIAamp Viral RNA Mini Kit (Qiagen, Hilden, Germany). cDNA was synthesized as per the protocol described by the manufacturer (Fermentas, Hanover, USA) using either oligo dT, random hexamer or SARS-CoV-2-specific primers as per the requirement. qRT-PCR was carried out to amplify SARS-CoV-2 “N” gene (Forward primer: 5’-ATACAATGTAACACAAGCTTTC-3’b and reverse primer: 5’-AGCAAAATGACTTGATCTTTG-3’) in a 20 μl reaction mixture containing gene-specific primers, template, and iTaq™ Universal SYBR® Green Supermix (Bio-Rad, USA) and has been previously by our group (23).

### Quantitation of m6A methylome in SARS-CoV-2 infected cells

Quantification of m6A modified RNA was carried out by using EpiQuikTM m6A RNA Methylation Quantification Kit (Colorimetric) (EpiGentek, Farmingdale, NY) according to the manufacturer’s protocol. Briefly, the RNA was isolated by TRI reagent (Sigma–Aldrich Steinheim, Germany). The isolated RNA along with negative and positive controls (supplied with the kit) were diluted in TE buffer and allowed to bind in the 8-Well Assay Strips in presence of binding solution. The RNA bound to the individual well of Assay Strips was detected using capturing (primary) and detecting (secondary) antibodies. The optical densities (OD) were taken at 450 nm in the microplate reader (Multiskan GO, Thermo Fisher Scientific, Helsinki, Finland). The OD values were normalized with negative controls and the absolute amount of m6A modified RNA (%) was calculated by comparing with the positive control.

### CHIP assay

CHIP assay was carried out to evaluate the interaction of viral RNA/mRNA with cellular proteins viz. cap-binding protein (eIF4E) and hnRNPA1 as per the previously described method (65, 66) with some modifications. Briefly, Vero cells, in triplicates were infected with SARS-CoV-2 at MOI (multiplicity of infection) of 5. At indicated time post-infection, the cells were treated with 1% formaldehyde for 10 min to covalently cross-link interacting proteins and nucleic acid. Thereafter, the cross-linking reaction was stopped by the addition of 125 mM glycine followed by washing the cells with ice-cold PBS. The cells lysates were prepared in immunoprecipitation (IP) buffer [150 mM NaCl, 50 mM Tris-HCl (pH 7.5), 5 mM EDTA, 0.5% NP-40, 1% Triton X-100 plus protease and phosphatase inhibitor cocktail] and sonicated in a Qsonica Sonicator Q500 (Qsonica, Newtown, CT, USA)(6 pulse of 15 sec at an amplitude of 40%). The sonicated cell lysate was centrifuged for 10 min at 12,000g. The clarified cell lysate was mixed with 10 units of RiboLock RNase Inhibitor (Thermo Scientific, USA) and then incubated with primary antibody or an equivalent volume of IP buffer (beads control) for 45 minutes at room temperature. Thereafter, 40 μl (5 ng/μl) of Protein A Sepharose® slurry, prepared as per the instruction of the manufacturer (Abcam, USA) was added into each reaction and incubated overnight at 4°C on a rotary platform. The beads were then washed 5 times in the IP buffer. In order to reverse the cross-linking, the complexes were incubated with proteinase K (20 mg/ml final concentration) at 56°C for 40 minutes. Finally, the reaction mixtures were centrifuged at 2000 g for 1 min and the purified RNA (TRI reagent) from the supernatant was subjected to cDNA preparation and quantitation of SARS-CoV-2 RNA (N gene) by qRT-PCR.

### 5’ cap synthesis assay

To evaluate the effect of DZNep on the synthesis of the 5’ cap of viral mRNA, a cell free interaction assay between viral mRNA and purified eIF4E protein was performed as described previously (67–69). Briefly, uninfected Vero cell lysate prepared with non-denaturing agents was incubated with Protein A Sepharose beads bound to α-peIF4E. The unbound α-peIF4E was removed by washing with IP buffer and stored at −70°C in a deep freezer. Next, Vero cells, in triplicates were infected with SARS-CoV-2 at MOI of 5. DZNep or vehicle control was added at 3 hpi. Subsequently, cells were lysed at 8 hpi and subjected to RNA extraction using TRI reagent. An equal amount of viral RNA (normalized by qRT-PCR) was then incubated with the complex containing peIF4E-αpeIF4E-Protein A Sepharose complex for 30 minutes in immunoprecipitation (IP) buffer [150 mM NaCl, 50 mM Tris-HCl (pH 7.5), 5 mM EDTA, 1% Triton X-100 plus RNase, protease and phosphatase inhibitor cocktail]. The complex was then subjected to centrifugation at 2000g for 2 minutes and washed 5 times with PBS. The pellet was subjected to RNA extraction using TRI reagent, followed by cDNA preparation (using oligo dT) and quantitation of SARS-CoV-2 mRNA (N gene) by qRT-PCR.

### Cross linking immunoprecipitation (CLIP) assay

Cross-linking immunoprecipitation was conducted as described previously with some modifications (70–73). Confluent monolayers of Vero cells were infected with SARS-CoV-2 at MOI of 5 for 1h in the presence of DZNep or vehicle control, followed by washing with PBS and the addition of fresh MEM having either DZNep or vehicle control. Cells were then treated at 2 hpi with 1% formaldehyde for 10 min to covalently cross-link the interacting proteins and nucleic acid. Thereafter, the cross-linking reaction was stopped by addition of 125 mM glycine (final concentration), followed by washing the cells with ice-cold PBS. The cells lysates were prepared by incubating cells with 1 ml of immunoprecipitation (IP) buffer [150 mM NaCl, 50 mM Tris-HCl (pH 7.5), 5 mM EDTA, 1% NP-40 plus protease and phosphatase inhibitor cocktail] for 10 minutes. Thereafter, the cell lysate was subjected to low centrifugation at 1000 g for 10 minutes. The top 1/3^rd^ portion (~300 μl), called the cytosolic fraction, was collected in a fresh tube and clarified again by centrifugation at 6000 g for 10 minutes (74). The pellet was discarded and the clarified cytosolic fraction was mixed with 10 units of RiboLock RNase, protease and phosphatase inhibitor cocktail and incubated with the Protein A Sepharose-bound m6A specific primary antibody (reactive antibody), Protein A Sepharose bound-phospho ERK antibody (non-reactive antibody) or an equivalent volume of IP buffer (beads control) overnight at 4°C on a rotary platform. Finally, the reaction mixtures were centrifuged at 2000g for 1 min and washed five times with PBS. The precipitated protein-RNA complex was resuspended in 100 μl PBS for quantification of the m6A-bound proteins (Western blot analysis).

### siRNA knockdown

Vero cells, in triplicates, were grown at ~75% confluency in 6 well plates and transfected with target- or negative control siRNAs **(Table S2)** using Lipofectamine 3000 as per the manufacturer’s (Invitrogen, Carlsbad, USA) instruction. At 48 h post-transfection, cells were infected with SARS-CoV-2 at MOI of 1 and the virus released in the infected cell culture supernatant at 24 hpi was quantified by plaque assay. The cell pellet was subjected to Western blot analysis to probe the respective cellular protein.

### Statistical analysis

Pairwise statistical comparisons were performed by two-tailed Student’s t-test in GraphPad Prism 8 software.

## DATA AVAILABILITY

Complete genome sequence of SARS-CoV-2 used in this study is available in GenBank with Accession number MW555598. The virus has also been deposited in the microbial repository at National Centre for Veterinary Type Cultures, Hisar, Haryana (India) with Accession Number VTCCAVA294.

## ACKNOWLEDGEMENTS

This work was supported by Indian Council of Agricultural Research, New Delhi (grant number IXX14586 to N-Ku and NASF/ABA-8027/2020-21 to N-Ku and B.R.G.) and Science and Engineering Research Board, Department of Science and Technology, Government of India (grant number CVD/2020/000103 to N-Ku).The funders had no role in study design, data collection and analysis, decision to publish, or preparation of the manuscript.

